# Targeting Autophagy Accelerates Intestinal Repair after Acute Ionizing Radiation

**DOI:** 10.64898/2026.07.06.736694

**Authors:** Madhuri Chaurasia, Aayushi Singh, Krishnamurthy Natarajan, Kulbhushan Sharma

## Abstract

Radiation exposure induces systemic and cellular damage, contributing to acute radiation syndrome and long-term effects such as premature aging and carcinogenesis. At the cellular level, radiation triggers apoptosis, mutation, and transformation through oxidative damage and activation of pathways including ER stress–mediated autophagy. Autophagy plays a context-dependent dual role in stressed cells, but its contribution to intestinal recovery after acute radiation remains unclear. Here, we evaluated combinatorial radiomodification using gamma radiation (8 Gy) and autophagy modulators in whole-body irradiated C57BL/6 mice (8–10 weeks old, n = 10). Mice were treated with autophagy inducers or inhibitors and euthanized at 3-, 8-, and 30-day post-irradiation. The jejunal–ileal region was analyzed via antioxidant assays, immunoblotting, H&E staining, and immunohistochemistry. Radiation significantly altered oxidative stress and autophagy markers, including increased LC3-II and decreased SQSTM1/p62. Autophagy induction enhanced intestinal proliferation (as measured by Ki-67), whereas inhibition impaired regeneration. Rapamycin pretreatment improved survival and reduced markers of intestinal injury following 8 Gy total body irradiation (TBI), whereas chloroquine exacerbated several injury-associated parameters. Overall, our findings suggest that targeted modulation of autophagy is a promising strategy for alleviating radiation-induced gastrointestinal injury and provide mechanistic insights relevant to therapeutic development.

## 1. Introduction

In the environment, humans are exposed to a variety of radiation, including UV rays, X-rays, and gamma rays, through medical interventions, radiation incidents, and accidents. Radiation exposure results in cellular instability due to the generation of reactive oxygen species (ROS) and reactive nitrogen species (RNS), which cause macromolecular damage at the cellular level and, ultimately, magnified organ damage, leading to both acute radiation syndrome and delayed effects. Hematopoietic, gastrointestinal, skin, and vascular endothelium are among the most radiosensitive organs (Amaravadi, R.K., et al., 2007; Apel, A., et al.,2008; Ayala, A., 2014).

Based on the dose of radiation exposure, acute radiation syndromes can be hematopoietic (HI) or Bone marrow syndrome, gastrointestinal (GI), and central nervous system (CNS) syndrome. Doses in the range of less than 7 Gy result in hematopoietic syndrome, which can be identified with an overall decline in blood cells (pancytopenia), increased susceptibility of radiation-exposed people to several infections, and hemorrhage. GI syndrome occurs following a whole-body exposure to more than 8 Gy (Bala, M., et al., 2015; Cao, C. et al., 2006).

Ionizing radiation causes macromolecular damage and metabolic imbalances, eliciting several intracellular responses that collectively determine the fate of the irradiated cell (Chaurasia, M. et al., 2016; Verma, N., and Tiku, A.B., 2022). Autophagy is one such process that can be elicited under numerous stress conditions, such as hypoxia, nutrient deficiency, and pathogenic infections (Chaurasia, M., et al., 2019; Chen, N. and J. Debnath, 2010). In cases of radiation and other traumatic situations, basal levels of autophagy are constitutively maintained within cells to maintain homeostasis. During stress, autophagy is modulated manifold to recycle damaged constituents.

Numerous efforts in autophagy and cancer biology have aimed to influence tumor survival. Only a few studies in the literature have suggested a role for autophagy in radiation exposure of normal cells or healthy individuals with no prior history of illness (Chen, Y., M.B. Azad, et al., 2009; Coleman, C.N. et al., 2004; Eke, I. et al., 2018). The important question is which therapeutic strategies should be employed by the clinicians to implement following radiation exposure to improve survival outcomes in exposed individuals, including first responders and army personnel. Specifically, what strategy do they use to modulate autophagy levels (induction or inhibition) to improve patient survival? Thus, to assess the effect of modulating systemic autophagy, the pharmacological autophagy modulators Rapamycin and Chloroquine were used in a mouse model. Rapamycin is an FDA-approved drug and has been used for several years to treat renal cell carcinoma and mantle cell lymphoma (Eke I, et al., 2018; Dainiak, N., 2002; Darzynkiewicz, Z. et al., 2008). It is an immunosuppressant that was originally used in transplant patients (Eke I et al., 2018). Rapamycin induces Macroautophagy through the inhibition of MTOR/ mechanistic target of rapamycin kinase complex 1 (MTORC1), which is the complex of MTOR with Raptor (Degenhardt, K. et al., 2006; Dorr, H. and V. Meineke, 2011; Gera, J. and A. Lichtenstein, 2011). On the other hand, Chloroquine is an endosomal acidification inhibitor that inhibits autophagy at later stages by raising lysosomal pH, thereby preventing the fusion of autophagosomes with lysosomes for cargo clearance. It is traditionally used as an antimalarial drug and has recently emerged as an anti-cancer agent and a chemosensitizer when used in combination with other anticancer drugs (Homewood, C.A. et al., 1972). It has been shown to inhibit cell growth and induce cell death in various cancer types.

In the present study, we have shown the role of prophylactic modulation of autophagy in the recovery of radiation-induced gastrointestinal damage in whole-body-irradiated C57BL/6 mice.

## 2. Materials and methods

### 2.1 Animals

Female C57BL/6 mice (8–10 weeks old) were used for all experiments. Animals were sourced from an accredited institutional facility and maintained under pathogen-free conditions, with routine health monitoring, at standard mouse housing conditions: 23–25 °C, 55 ± 5% relative humidity, and a 12 h light and 12 h dark cycle. Mice were fed standard rodent feed (bought from Golden Feeds, India). All animals were healthy and had no prior experimental procedures. Mice were grouped and injected with autophagy modulators: Chloroquine (Sigma-Aldrich, C6628) or Rapamycin (Sigma-Aldrich, PZ0020). Chloroquine was reconstituted in PBS and administered at a 10 mg/kg dose via the intraperitoneal (i.p.) route. Rapamycin was reconstituted in DMSO at 20 mg/ml and further diluted in PBS containing 5% DMSO to get the desired 2 mg/kg body weight dose, which was administered through the (i.p.) route. The autophagy modifiers were administered 1 h before irradiation, unless otherwise mentioned. Administration of the Vehicle alone (PBS containing 5% DMSO) did not induce any detectable toxicity. For the endpoint studies and biochemical assays, the mice were euthanized by CO_2_ asphyxiation followed by cervical dislocation. All proper controls were included in the study.

### 2.2 Irradiation Procedure

For in vivo experiments, at least 10 mice/group were irradiated with the Tele-Cobalt Facility, Bhabhatron II (Panacea Medical), using a field size of 35 cm x 35 cm at an SSD of 80 cm in the Irradiation Center. Mice in each group were given 1 h prior to treatment of autophagy modulators and exposed to whole-body radiation of 8 Gy from a ^60^Co γ -ray irradiator having a dose rate of 1.25 Gy/min, followed up for survival till 30 days. For western blotting and other assays, three mice from each group were sacrificed on the third, eighth, and thirtieth day after irradiation. All experiments complied with the Institutional regulations on animal welfare protocols and were approved by the Institute’s ethics committee on laboratory animals.

### 2.3 Survival Study in Mice

For a survival study, mice were randomly assigned to six groups (n = 10 per group) using a computer-generated randomization sequence in Microsoft Excel. Randomization was performed prior to the start of the experiment, with animals from different cages distributed across groups to minimize cage-related confounding. Group allocation was generated by the investigator prior to the start of the experiment. During the experiment, investigators were aware of the group allocation because of the treatments administered. However, outcome assessment and data analysis were performed with investigators blinded to group allocation to minimize bias. Experimental groups were: unirradiated Control, IR-only radiation, Rap, CQ, Rap+IR, and CQ+IR. Mice in each group received 1 h of prior treatment with autophagy modulators, were exposed to 8 Gy of whole-body radiation, and were assessed for survival for 30 days. Animals were monitored daily for clinical signs of distress, including weight loss, hunched posture, diarrhea, inability to stand, and reduced activity.

### 2.4 Western blot analysis

Jejunal and ileal sections from mice were used to isolate intestinal cells (n = 4). The Intestine was flushed with ice-cold PBS, and intestinal cells were isolated by overnight collagenase treatment at 4 °C, followed by homogenization using an MP homogenizer in RIPA buffer with Protease and phosphatase inhibitor cocktail (Thermo Fisher, Pierce Protease Inhibitor Mini Tablets, 88665) on ice. Cell lysates were centrifuged at 4°C (14000 rpm, 20 min), and the protein supernatant was transferred to new microcentrifuge tubes. Protein concentrations in the samples were determined using the BCA Protein Assay Kit (Pierce, Rockford, IL, USA). 20 µg of total protein was resolved by 6%, 12%, or 15% SDS-PAGE, followed by protein transfer to PVDF membranes. The membranes were blocked in PBS containing 0.1% Tween-20 (PBST) with 3% bovine serum albumin at room temperature for 1 h. To detect multiple proteins on the same membrane, the blot was cut horizontally into strips based on the approximate molecular weights of the target proteins, and each strip was probed with the corresponding primary antibodies. The following primary antibodies were used: SQSTM1/p62 (Sigma-Aldrich, P0067), PARP1 (Pierce, MA5-15031), ACTB (SC-47778), GAPDH (SC-25778), and LC3A/B (CST-4108). Secondary antibodies were used from Santa Cruz. Blots were detected using ECL reagents (Amersham Pharmacia Biotechnology, Buckinghamshire, UK). Only the blots having band intensities within the linear range were processed for quantitation. The blots were quantified using ImageJ software.

### 2.5 Histology and immunohistochemical (IHC) staining for Ki-67

GI tissues (ileal and jejunal sections) were fixed in 10% neutral-buffered formalin (SRL) and embedded in paraffin. GI samples were sectioned at 5 μm thickness using a Leica microtome and stained with H&E (hematoxylin and eosin). The sections were scored for Surviving Villi Number and Villi Height. Only complete sections, which included the opening of the crypt and the full length of villi from base to the tip, were considered for scoring. All counts and measurements from each tissue specimen were obtained “blind” from a minimum of 3 coded sections. ∼5–7 fields of view (FOV) per mouse/section were assessed.

### 2.6 Fluorescence-based Immunohistochemistry (IHC-F) in intestinal sections

To assess intestinal crypt proliferation, Ki-67 fluorescence-based IHC was performed on jejunal sections from mice according to the manufacturer’s protocol. Briefly, sections embedded in paraffin were deparaffinized, rehydrated, and incubated with blocking buffer for 30 minutes, followed by incubation with Ki-67-FITC monoclonal primary antibody (BioLegend, 652409) at a 1:800 dilution overnight at 4°C. Corresponding tissue sections without primary antibody served as negative controls. Sections were examined by fluorescence microscopy to capture FITC and DAPI-stained images using an Olympus (IX51) microscope, which were acquired at 10X and 40X magnification for quantification and are presented in the results. Ki-67 analysis was focused on the stromal compartment. For Microtubule-associated protein 1 light chain (LC3-II) staining, sections were deparaffinized, antigen retrieved in citrate buffer (pH 6.0; Dako, Carpinteria, CA), and endogenous peroxidase quenched. Sections were incubated in blocking buffer (0.1% bovine serum albumin in PBS) before exposure to the LC3A/B antibody (recognizes both LC3-I and LC3-II; however, punctate cytoplasmic staining primarily reflects LC3-II localized to autophagosomal membranes). Increased LC3 staining was interpreted as evidence of autophagosome accumulation. Tissue sections were incubated overnight with anti-LC3A/B antibody (CST-4108; dilution 1:100) at 4 °C. After necessary washing steps, sections were incubated with goat anti-rabbit HRP-conjugated secondary antibody. SuperPicture TM 3rd Gen IHC detection kit (87-9673; Invitrogen) was used for signal detection and color development. All the IHC slides were mounted and visualized under a bright-field microscope, and images were captured at microscopic magnification (20X magnification). To determine staining specificity, appropriate controls were run in parallel with the experimental sections.

### 2.7 Lipid peroxidation assay

Lipid peroxidation was estimated spectrophotometrically by the modified thiobarbituric acid-reactive substance (TBARS) method (as described by Varshney and Kale, 1990). Lipid peroxidation was measured in the GI tissues of mice harvested at days 3 and 8 post-treatment (n = 6). Cell lysate / GI tissue lysate (80 µl) was mixed with 580 µl LP Buffer (0.15 M KCl + 10 mM Tris-HCl, pH 7.4), to which 30% TCA (166 µl) was added and vortexed. Then, 52 mM (166 µl) TBA was added. The tubes were placed in a water bath in the dark for 45 min at 80 °C, cooled on ice, and centrifuged at room temperature for 10 min at 3,000 rpm. The absorbance of clear supernatant was measured against a reference blank at 532 nm in a spectrophotometer. The amount of MDA formed in a sample was estimated according to the equation:

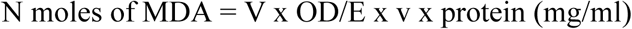

where, V = final volume of test solution (ml), OD = optical density, E= extinction coefficient and v = sample volume.

Finally, the end product of jejunal lipid peroxidation (MDA) was expressed in nmol/mg tissue.

### 2.8 Biochemical Measurements

#### SOD assay

Activities of the antioxidant enzyme Superoxide dismutase (SOD) were measured in mice GI tissues harvested 3 days post-treatment (n = 6). The SOD activity assay is based on the auto-oxidation of pyrogallol, a process that is highly dependent on superoxide, which is the substrate for SOD. The auto-oxidation of this compound is inhibited in the presence of SOD, whose activity was then indirectly assayed at 420 nm according to the method of Marklund and Marklund, 1974. The results were represented as SOD units/mg.

#### GSH assay

The level of the non-enzymatic cellular antioxidant glutathione was measured according to the method of Moron et al., 1979. The estimation was performed by measuring GSH activity in GI tissues harvested from mice at 3 days post-treatment (n = 6). A GSH stock solution (1 mM in 5% TCA) was prepared for generating the GSH standard curve by serial dilution in triplicate. The sulfhydryl group of GSH reacts with DTNB (5,5’-dithio-bis-2-nitrobenzoic acid) and produces a yellow -coloured product. The protocol is a colorimetry-based assay. Samples in which GSH levels were to be analyzed were assayed by taking 30 µL of the sample and 200 µL of DTNB, followed by incubation at 37°C in the dark. Activity was measured by taking absorbance at 415 nm using an automated microplate reader (Bio-Tek, Winooski, USA).

### 2.9 Terminal deoxynucleotidyl transferase dUTP nick end labeling (TUNEL) assay

TUNEL assay was performed on jejunal sections from mice using the in-situ death detection kit (Sigma-Aldrich, 11684795910-roche) as per the manufacturer’s instructions. Briefly, sections were deparaffinized, pretreated with proteinase K, and endogenous peroxidase was quenched using H_2_O_2_. The sections were incubated with FITC-tagged terminal deoxynucleotidyl transferase (TdT), and counterstained with DAPI. Stained sections were visualized under a Zeiss Axio Vision microscope. Representative images captured at 20X magnification are presented in the results. Apoptotic cells were scored in the jejunum crypts.

### 2.10 FACS analysis

Quantitative flow cytometric analysis was performed on mouse bone marrow cells isolated from the femora and tibiae of mice (n = 3). Cells were isolated by flushing bone marrow with 1-3 mL phosphate-buffered saline (without Mg2^+^ and Ca2^+^) supplemented with 5 mM EDTA plus 1% fetal bovine serum. Following incubation in RBC lysis buffer (150 mM ammonium chloride, 10 mM potassium bicarbonate, and 0.1 mM EDTA) on ice for 10 minutes to lyse red blood cells, the cells were spun down, resuspended in ice-cold PBS, and washed. The cell pellet was then fixed with 4% PFA for 15 min, followed by permeabilization with a permeabilization buffer (1X TBS + 0.1% Triton X-100). The cell pellet was resuspended in PBS and filtered through a 70-μm filter to remove aggregated cell clumps and debris. Cells were blocked and then stained with a Ki-67-FITC antibody for 1 h at room temperature in the dark. Flow cytometry analyses were carried out on the BD FACS Caliber Cell Analyzer (BD Biosciences). Briefly, debris and non-cellular events were excluded based on forward scatter (FSC) and side scatter (SSC) characteristics, and doublets were removed using FSC-A versus FSC-H gating. Ki-67-positive cells were identified by setting the positivity threshold based on an appropriate unstained control.

### 2.11 Statistical analysis

All experiments were performed at least three times and then analyzed unless otherwise stated. All data are presented as mean ± S.D. of the mean from triplicates. Statistical analysis was performed by one-way ANOVA using Tukey’s multiple comparison test or with two-way ANOVA with Tukey’s multiple comparison test using the Graphpad Prism software for Windows (Graphpad, Inc., California Corporation). P< 0.05 was considered statistically significant. For the mouse survival study, analyses were performed using Kaplan-Meier curves, and statistical significance was assessed using the log-rank (Mantel-Cox) test.

## 3. Results

### 3.1. Effect of autophagy modifiers on the survival of whole-body irradiated C57BL/6 mice

To explore the in vivo radiomodulative potential of pharmacological modifiers of autophagy, Rapamycin (Rap, 2 mg/kg body weight) and Chloroquine (CQ, 10 mg/kg body weight) were injected (n= 10) through the intraperitoneal route in C57BL/6 mice 1 h before total body γ-irradiation (TBI). Upon irradiation, various symptoms, including a gradual loss in body weight, food and water intake, and ruffled fur, were observed, and consequently, 50% of the mice died within 15 days of exposure (8 Gy IR dose is the LD50 dose in these mice) in radiation alone as compared to the unirradiated control mice (Figure 1). Treatment with Rap provided more than 50% protection in 8 Gy-irradiated mice as compared to the radiation control. In contrast to this, CQ+IR-treated mice indicated enhanced reduction in body weight, food and water intake, and with lesser survival as compared to IR alone (20 % less) (Figure 1A and B). Here, in this study, we observed a similar mouse survival trend to our previous finding: a significant survival advantage for irradiated mice that had received prior Rapamycin treatment as an autophagy inducer (Chaurasia, M. et al., 2019).

**Figure 1.**
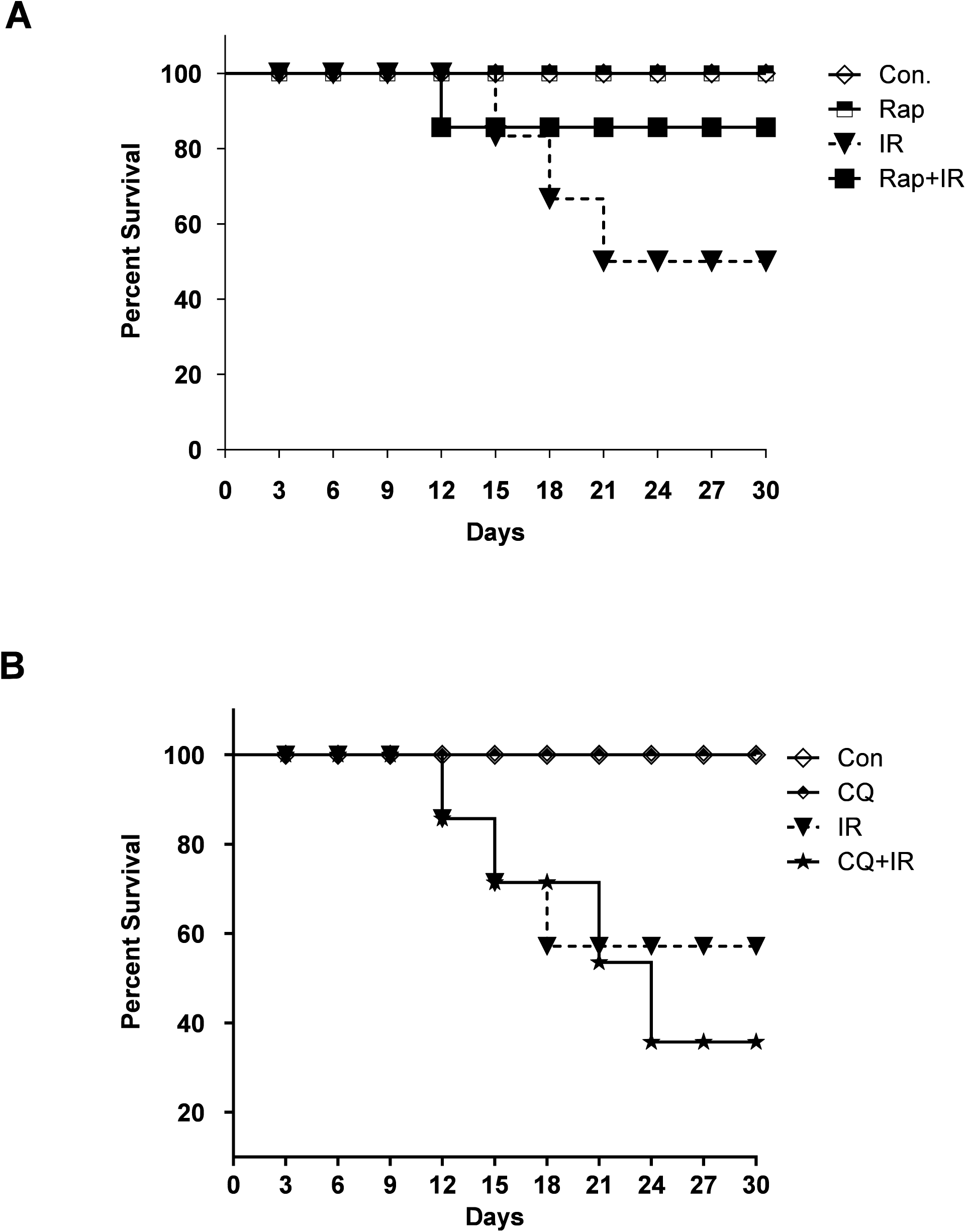
Radiation-induced autophagy is pro-survival under in vivo conditions. (A) The effects of the autophagy inducer, Rapamycin (2mg/kg body weight), on survival during the first 30 days after 8Gy irradiation in mice. C57BL/6 mice were randomized into four groups: control, IR, Rap., Rap.+IR. Rapamycin was administered i.p. as a single dose 1 h before irradiation. Mice were weighed, and lethality was scored daily for the first 30 days. Kaplan-Meier analysis was performed. Each treatment group contained at least ten animals. (B) The effect of the autophagy inhibitor Chloroquine (10 mg/kg body weight) on animal survival was studied for the first 30 days after 8 Gy irradiation in mice. C57BL/6 mice were randomized into four groups: control, IR, CQ alone, and CQ+IR. CQ was administered via intraperitoneal (i.p.) injection in a single dose, 1 h prior to irradiation. Mice were weighed, and lethality was scored daily for the first 30 days. Kaplan-Meier analysis was performed for mice receiving 8 Gy of total-body irradiation.

### 3.2. Effect of autophagy modifiers on irradiated C57BL/6 mice intestinal damage recovery

To explore radiation-induced intestinal changes in drug-treated mice, three mice per group were euthanized on days 3, 8, and 30 post-irradiation, and H&E staining was performed. The villi from the jejunum and ileum of healthy untreated controls were tall and cylindrical, with an adequate number of crypt cells having optimal length. In irradiated animals, villus height and cellularity were reduced (Figure 2A). At 72 h after irradiation, the number of crypts and villi, as well as villus height, were significantly decreased. These were associated with other gross histological changes, such as villi fusion and non-recoverable decreases in villi height (Figure 2A).

**Figure 2.**
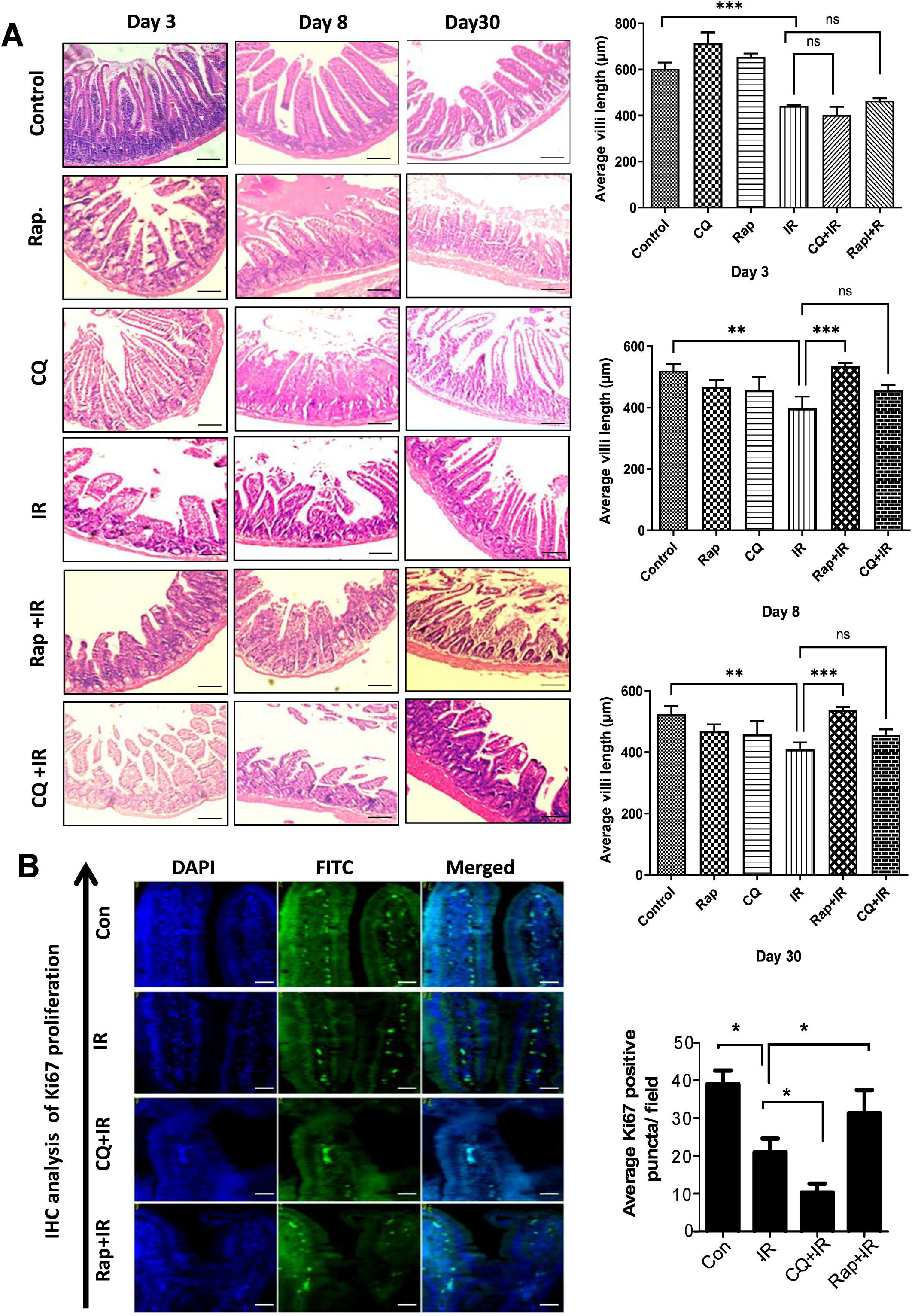
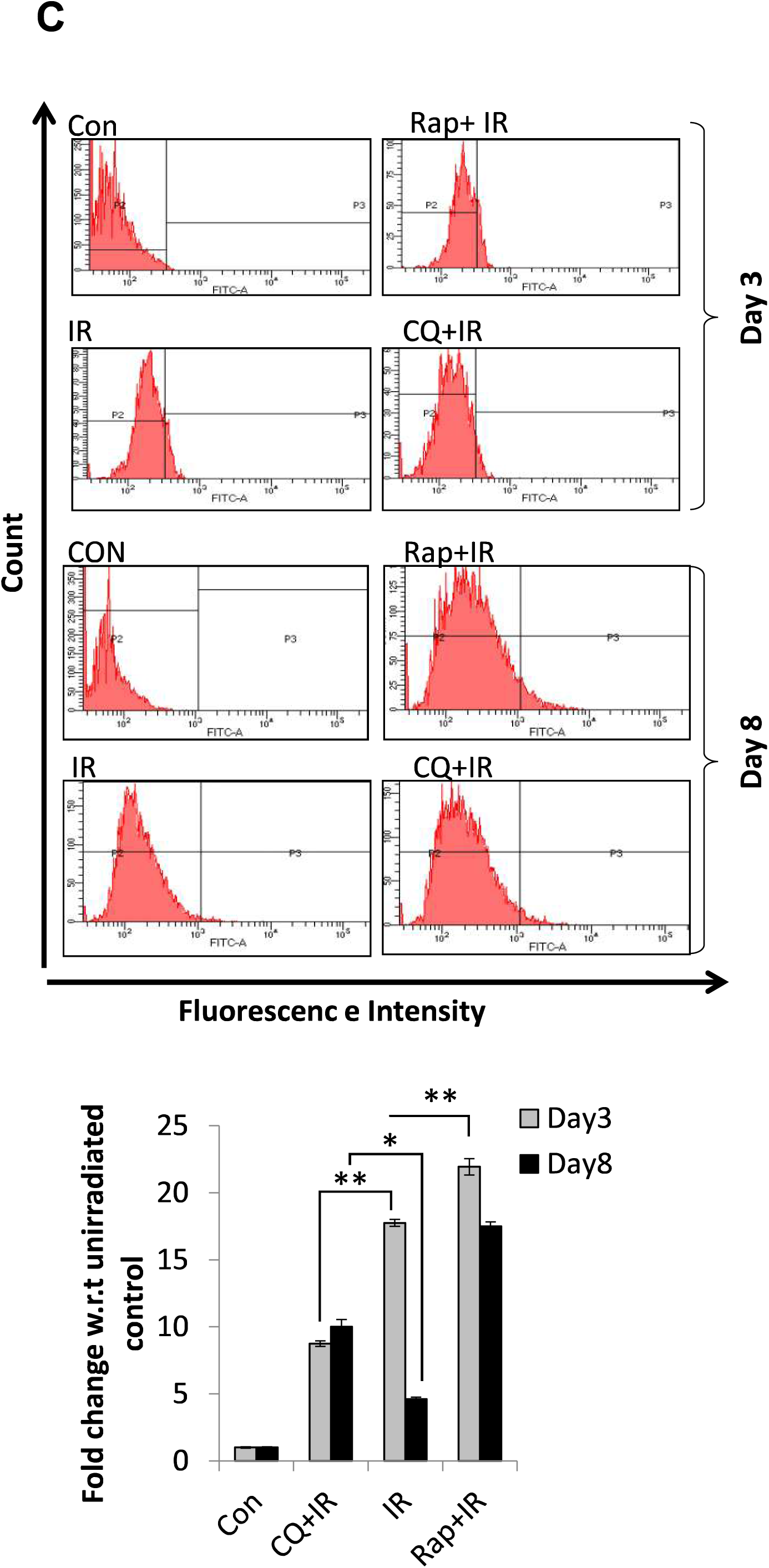
Effect of autophagy modifiers on the recovery of irradiated mice’s intestinal damage. **(A)** H&E-stained sections of C57BL/6 jejunum obtained at Day 3, Day 8, and Day 30, after exposure to whole-body irradiation (8 Gy), after single-dose administration of autophagy modulators (Rap or CQ) 1 h prior to irradiation. Representative micrographs were taken at 10X magnification (intestinal sections from six mice were examined in each group; n =6). Scale bar = 200 µm. **(B)** Fluorescence-based immunohistochemistry (IHC-F) analysis of Ki-67-positive cell proliferation in the presence of autophagy modifier drugs on day 8 post-exposure. Representative micrographs were taken at 40X magnification. Scale bar = 50 µm. **(C).** Quantitative analysis of Ki-67-positive cells indicates increased bone marrow cell proliferation in the presence of autophagy-modifying drugs on day 8 post-radiation exposure. Here, P2 and P3 correspond to Ki-67–negative and Ki-67–positive populations, respectively. Statistical analysis was performed using One-way ANOVA with Tukey’s multiple-comparison test. ns = non-significant, * p < 0.05, ** p < 0.01, and *** p < 0.001.

In Rapamycin-treated groups, there was no substantial additional reduction following irradiation, with villi length remaining significantly higher than in the IR-only group. In contrast, the CQ+IR group exhibited a pronounced decrease in villus height and cell numbers compared with the control, closely resembling the effects observed with irradiation alone, indicating that CQ did not confer protective effects, unlike Rapamycin.

Ki-67 is a widely used cell proliferation marker expressed in the nucleus of proliferating cells (Sobecki M, et al., 2016). Following radiation injury, increased Ki-67 expression in stromal cells reflects activation of the mesenchymal compartment, which plays a central role in epithelial regeneration. Proliferating stromal cells restore the intestinal stem cell niche by expanding the supportive mesenchyme, enhancing secretion of Wnt ligands, R-spondins, EGF, and HGF, and remodeling the extracellular matrix to facilitate crypt reconstitution (Kim WH, et al., 2023; Chen Y, et al., 2022). To study intestinal crypt proliferation profile upon radiation exposure, Fluorescence-based Immunohistochemistry (IHC-F) Ki-67 staining of intestinal sections was performed on days 3, 8, and 30 post-irradiation. IHC-F staining with Ki67-FITC antibody showed enhanced proliferation in rapamycin treated group, while less proliferation was observed in the autophagy inhibitor (chloroquine) treated groups (Figure 2B). Treatment with Rap 1 h before irradiation significantly counteracted radiation-induced early histological changes (within 72 h), and this effect continued to improve over time. On the other hand, CQ treatment prior to irradiation decreased the number of Ki-67-positive cells compared to the irradiated control. No alteration in the number of Ki-67 positive cells was observed in Rap/CQ alone when compared with the control group (Figure S1, S2). Radiation exposure-induced hematopoietic injury results in a rapid decline in hematopoietic progenitor cells, leading to pancytopenia (Dorr, H. and V. Meineke, 2011; Fliedner, T.M., et al., 2007; Galluzzi, L., et al., 2015; Ghosh, S.P., et al., 2012). Mice exposed to this level of total body irradiation displayed a rapid decline of all mature blood cell types, with a corresponding loss of hematopoietic progenitors in the bone marrow compartment. A greater recovery of bone marrow cells was observed with Rap treatment, suggesting that the induction of autophagy may aid hematopoietic recovery (Figure 2C).

### 3.3 Antioxidant status in murine GI tissue lysate

Highly reactive free radicals, ROS and RNS, generated after radiation exposure cause macromolecular damage in GI cells, and are important contributors to acute GI injury (Kim, Y.J., et al., 2012; Hall EJ and Giaccia AJ. 2006; Wu W, et al., 2025). To assess the contribution of autophagy-mediated reduction in radiation-induced damage, the activities of the antioxidant enzymes superoxide dismutase (SOD) and glutathione peroxidase (GPx), non-enzymatic GSH, and lipid peroxidation were assessed. Radiation caused a significant reduction in SOD (*p* < 0.05) activity, while GSH levels were marginally affected; Rap treatment was able to restore it remarkably (*p* < 0.01) (for both GSH and SOD) in the GI of radiation-exposed mice (Figure 3A and B). In contrast, CQ+IR treatment further reduced GSH levels and SOD activity compared with radiation-control mice (Figure 3A and 3B). Ionizing radiation causes toxicity and multiple forms of damage to vital biomolecules, either directly through the deposition of energy or indirectly via the decomposition of water molecules in the human body, which, in turn, leads to the generation of ROS such as hydroxyl, hydrogen peroxide, superoxide radicals, and RNS (Kim, Y.C. et al., 2014). These ROS and RNS react with lipids and other molecules present in cell and organelle membranes, initiating a chain reaction, i.e., lipid peroxidation (LPO), by abstracting additional hydrogen atoms. Thus, the formed LPO initiates the peroxidation of polyunsaturated fatty acids, leading to the production of Malondialdehyde (MDA) within stressed cells (Hacker, G., 2000). Thus, the formed LPO adduct alters membrane integrity, permeability, fluidity, and function of membrane-bound enzymes (Klionsky, D.J., et al., 2021; Lang, S.A. et al., 2007). Enhanced lipid peroxidation was observed in irradiated animals’ GI as compared to the unirradiated control group (p<0.05). Animals given the combined treatment of CQ+IR showed further enhanced levels of MDA formation (p<0.05). Significantly low levels of LPO adduct were formed in Rap+IR treatment as compared to the radiation control (p<0.01 for day 3 and 8) (Figure 3C).

**Figure 3.**
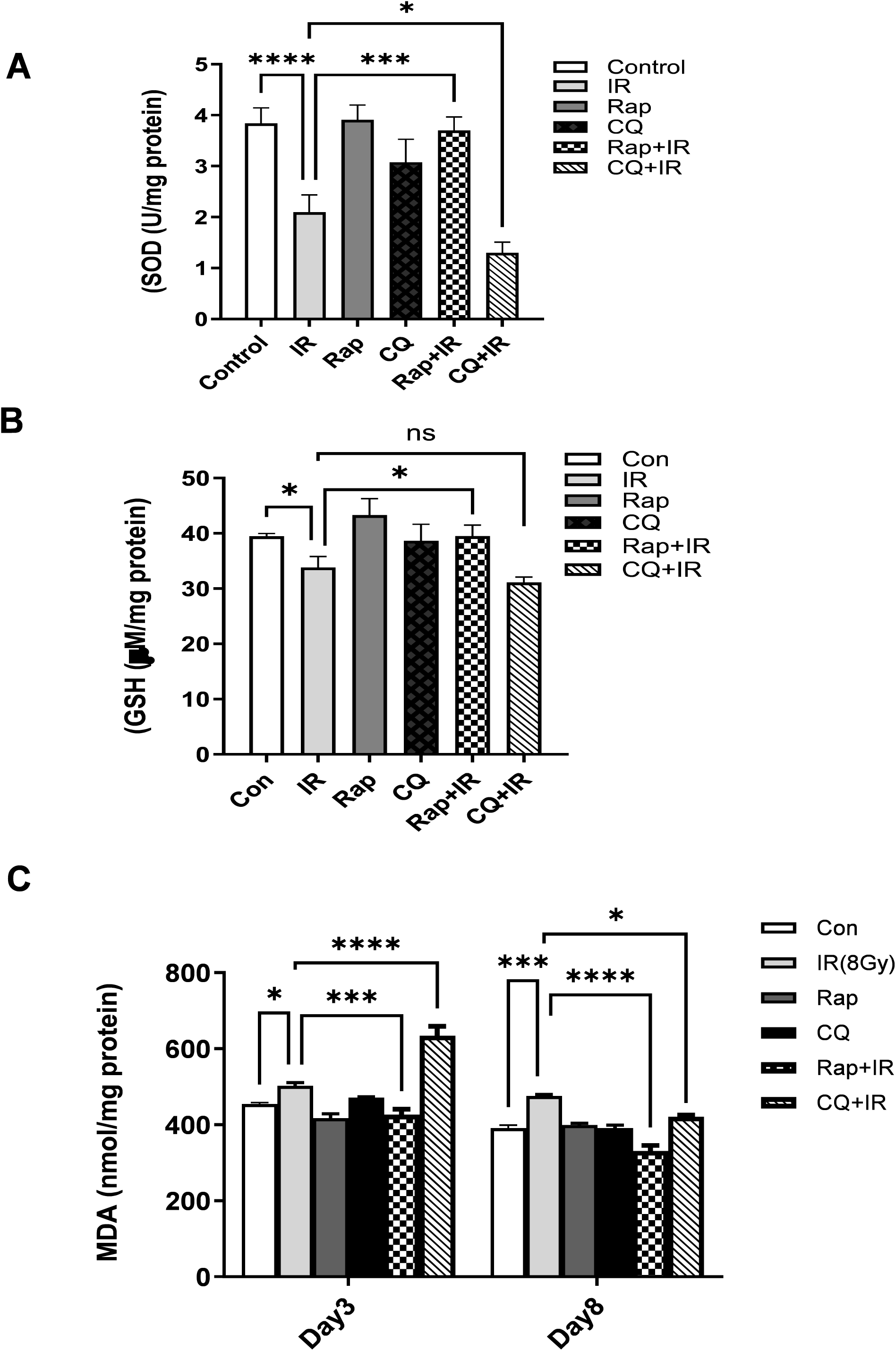
Antioxidant assays and lipid peroxidation in GI tissue lysates. **(A)** SOD activity level in the GI tissues of mice on day 3 post radiation exposure. **(B)** GSH level in the GI tissue of mice at day 3 post-treatment. **(C)** Lipid peroxidation in the GI tissue of mice on day 3 post-treatment. Induction of LPO in radiation-exposed C57BL/6 mice in the presence of autophagy modulators on day 3 and day 8 post-exposure. GI ileal-jejunal sections were harvested and used for LPO assay to determine MDA levels per mg protein. For A-B, Statistical analysis was performed using a One-Way ANOVA with Tukey’s multiple-comparison test. For (C), analysis was performed using a two-way ANOVA with Tukey’s post hoc test. * p < 0.05, ** p < 0.01, *** p < 0.001, and **** p < 0.0001.

### 3.4 Autophagic status in murine GI tissue

The radioprotective effect of Rap treatment in the mice’s intestines was evaluated on days 3 and 8 post-radiation exposure. IHC with the LC3-A/ B antibody showed increased LC3 levels (specifically corresponding to increased LC3-II (lipidated LC3), thus indicating high autophagic activity in irradiated animals. Rap +IR-treated mice showed a relatively greater accumulation of LC3-II-positive cells in GI tissue than IR- or CQ+IR-treated mice. Drug-alone groups didn’t show comparative LC3-II positive DAB staining. (Figure 4A). Further, immunoblot studies have clearly shown the effect of Rap and CQ pretreatment on LC3-II lipidation and p62 expression in the jejunum of lethally irradiated mice. LC3-II levels were significantly elevated (>2-fold; p < 0.05) at 72 h after 8 Gy exposure in the IR-treated group compared with the unirradiated control. A further boost of LC3 lipidation and a decline in p62 expression were observed in Rap+IR-treated groups as compared to IR alone (Figure 4B). In contrast, a reduction in LC3-II and an increase in p62 induction were observed in the CQ+IR-treated group as compared to IR (Figure 4B). These findings indicate modulation of autophagy markers in radiation-treated groups in the presence of autophagy modifiers. Although the absence of flux assays prevents distinction between increased autophagosome formation and impaired degradation.

**Figure 4.**
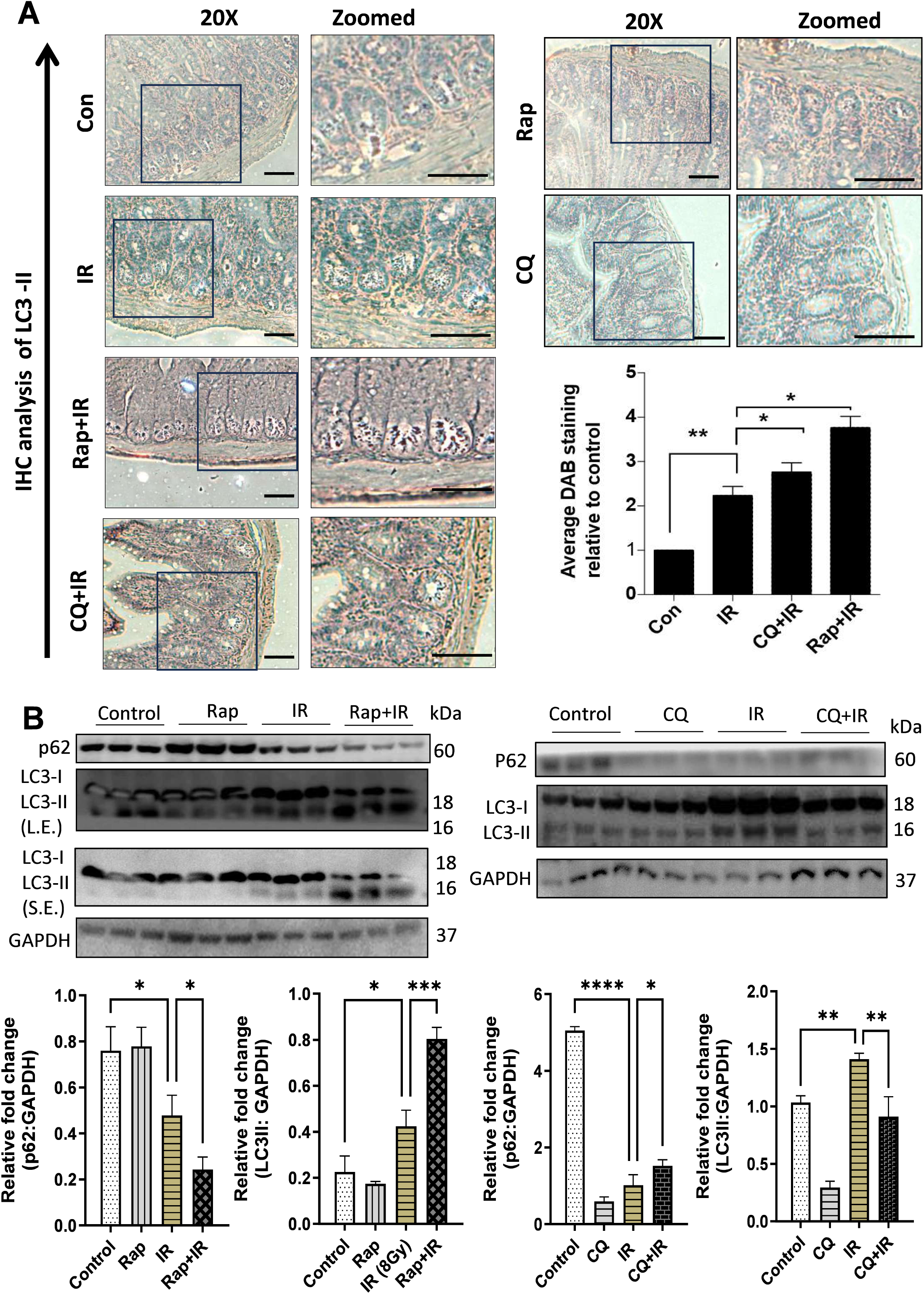
Autophagic status in mice GI tissue. **(A)** Autophagy status of mouse jejunal tissue using the LC3-A/B antibody by IHC marks: increased LC3 staining, specifically punctated staining, corresponds to enhanced LC3-II puncta accumulation. It indicates autophagosome accumulation, but does not independently measure autophagic flux. The effects of Rap and CQ treatments were evaluated on day 3 post-exposure. Representative images were captured at 20X. Scale bars = 100 µm and 200 µm for inset zoom-ins. **(B)** Western blot analysis of LC3-II and p62 levels in the mouse jejunal tissue lysate. The effects of Rap and CQ treatments were evaluated on day 3 post-irradiation. The bands were quantified by normalizing band intensities relative to GAPDH. S.E. short exposure, L.E. long exposure. Statistical analysis was performed using One-way ANOVA with Tukey’s multiple-comparison test. * p < 0.05, ** p < 0.01, *** p < 0.001 and **** p < 0.0001.

### 3.5 Programmed cell death studies in autophagy modifier-treated murine GI tissue

The relationship between radiation exposure and apoptosis has been widely established. Apoptosis is a programmed cell death phenomenon in which dead cells do not release toxic components to their surrounding microenvironment (Kim, Y.C., et al., 2014; Laplante, M. and D.M. Sabatini, 2013; Levine, B., 2007). The apoptotic process involves cleavage of cellular substrates via Caspase activation, DNA fragmentation, nuclear and cytoplasmic shrinkage, and membrane blebbing (Levine, B. and G. Kroemer, 2008). Autophagy and apoptosis exhibit a complex relationship depending on the type and extent of the stress or stimulus. Recent research has observed that radiation exposure is causally related to autophagy induction (Chaurasia, M., et al., 2019; Chen, Y., et al., 2009). Therefore, it would be worthwhile to study the relationship between these two processes (autophagy and apoptosis) in radiation-exposed conditions. To evaluate the intestinal epithelial cell apoptosis, the Terminal deoxynucleotidyl transferase dUTP Nick End Labeling (TUNEL) assay was performed (Figure 5A). The TUNEL assay substrate, i.e., terminal deoxynucleotidyl transferase (TdT)-mediated addition of labeled deoxyuridine triphosphate (dUTP) nucleotides to the 3’-OH ends of DNA strand breaks (generated in the later stages of apoptosis), aids in measuring the extent of apoptosis in those cells (MacNaughton, W.K., 2000). In our TUNEL studies, compared with the control, intestinal sections from irradiated mice showed a significant increase in TUNEL-positive apoptotic cells 3 days post-exposure. In Rap-pretreated groups, the number of apoptotic cells was markedly decreased compared with the radiation-treated group, whereas the opposite was observed in CQ+IR-treated animals. No significant alteration in the number of TUNEL-positive cells was observed in Rap/CQ alone-treated mice throughout the GI mucosa when compared with the control group. TUNEL-positive cells were observed throughout the GI mucosa of IR-treated mice on day 3 post-IR exposure. CQ+IR treatment significantly increased the number of TUNEL-positive epithelial cells in the small intestine of these mice (Figure 5A).

**Figure 5.**
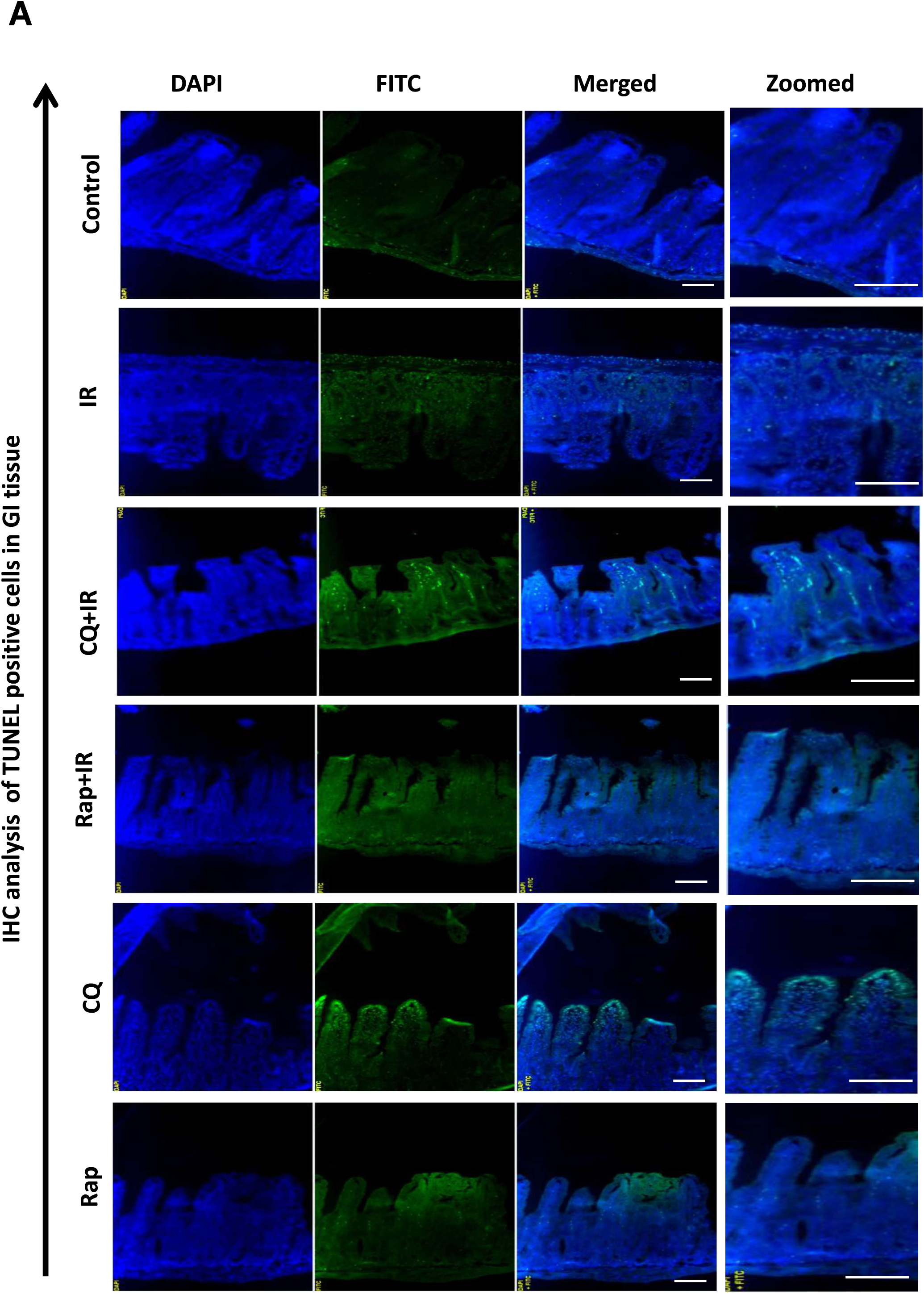
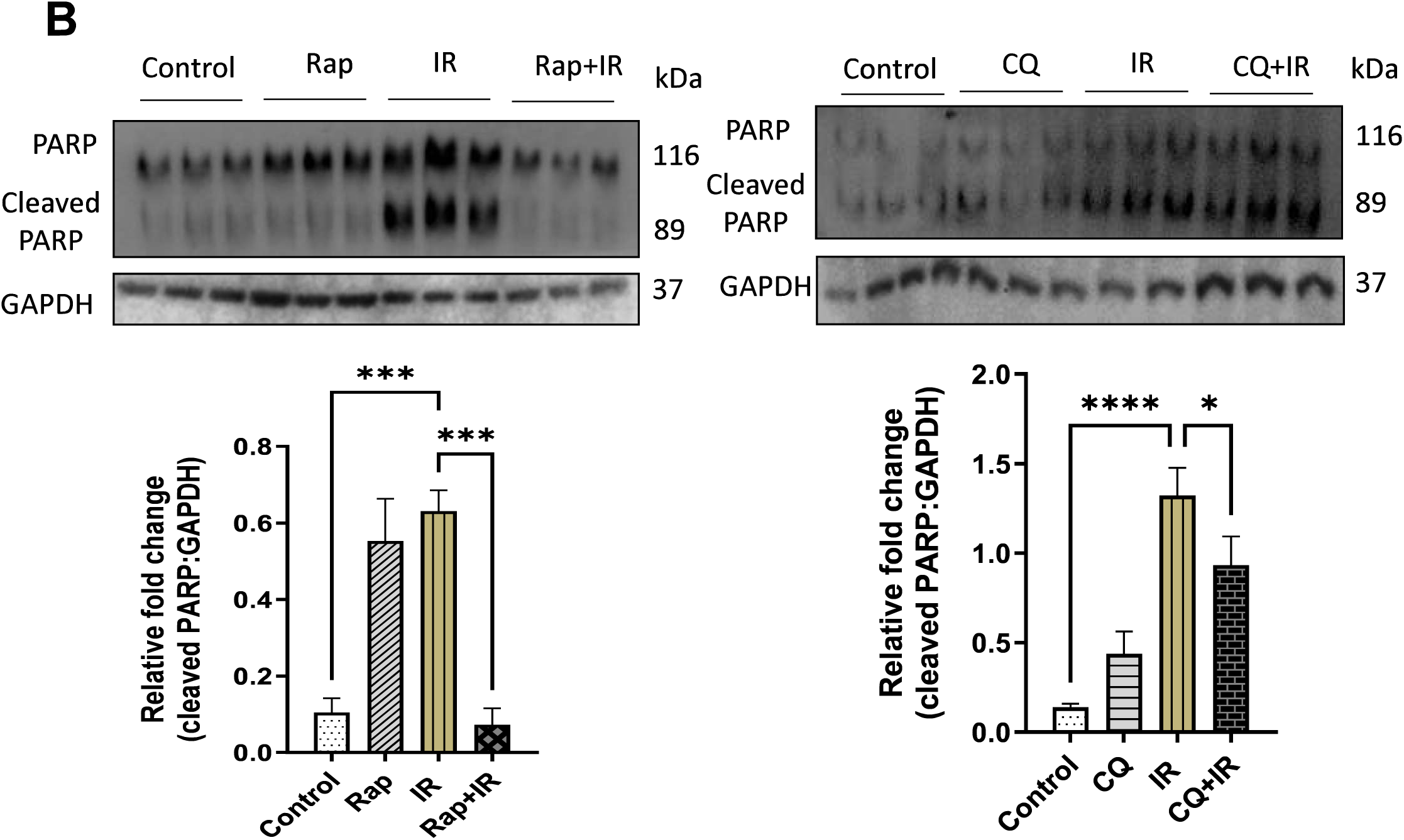
Programmed cell death studies in autophagy-modifier-treated murine GI tissue. **(A)** IHC analysis of TUNEL-positive cells in GI tissue on day 3 post-exposure. Stained sections were visualized under an upright fluorescence microscope; images were captured at 20X magnification to quantify TUNEL-positive cells per crypt. Images were quantified using ImageJ software. Scale bars: 100 µm and 200 µm for inset zoom-ins. **(B)** Western blot analysis of GI tissue lysate obtained at Day 3 post-irradiation from Rap+IR and CQ+IR treated mice tissue. Blots were probed with the intrinsic apoptosis marker, cleaved PARP; GAPDH was used as a loading control. Fold change was analyzed from biological triplicates; error bars (mean ± SD). One-way ANOVA with Tukey’s multiple comparisons. * p < 0.05, *** p < 0.001 and **** p < 0.0001.

We examined the effect of Rap pretreatment on cleaved PARP expression in the jejunum of lethally irradiated mice. Cleaved PARP expression was significantly increased at 72 h after IR exposure. The radioprotective effect of Rap treatment in the mice’s intestines was evaluated at day 3 post-exposure. Interestingly, Rap administration resulted in the suppression of cleaved PARP expression. However, Rap alone did not induce any significant change in cleaved PARP at both time intervals. In contrast, enhanced apoptosis was observed in CQ+IR-treated mice at day 3 post-exposure, as evidenced by increased cleaved PARP expression (Figure 5B). To summarize, both the TUNEL assay and Western blotting indicated reduced apoptosis in the Rap+IR-treated group, while induction of apoptotic markers was observed in the CQ+IR-treated groups.

## 4. Discussion

The growing incidence of possible exposure of humans to radiation has raised the need to develop countermeasures against radiation injuries. Radiation exposure causes toxicity and damages vital macromolecules such as nucleic acids, proteins, and lipids, etc., present in the cell (Amaravadi, R.K., et al., 2007; Apel, A. et al., 2008). The resulting effects of radiation exposure are dose-dependent. The hematopoietic syndrome can be subsequently managed with growth factors, bone marrow transplantation, blood transfusion, etc. (Measey, T.J. et al., 2018), but to date, there is no approved medical countermeasure to treat radiation-induced GI damage. Only a handful of studies are available on medical countermeasures for recovery from IR-induced GI damage (Gera, J. and A. Lichtenstein, 2011; Meistrich, M.L. and M. Kangasniemi, 1997).

Autophagy, a context-dependent phenomenon, is activated under stress conditions to promote cell survival or to sensitize cells (Chaurasia, M. et al., 2016; Meng, L.H. and X.F. Zheng, 2015). It can have opposing effects on tumorigenesis, i.e., both promoting and inhibiting tumor growth (Njie-Mbye, Y.F. et al., 2013). Treatment strategies, including autophagy induction, have been successful in treating human gastric adenocarcinomas and hepatocellular carcinoma. MTOR expression has been shown to increase, and tumor growth and angiogenesis are reduced in experimental models following rapamycin treatment (White, E. and R.S. DiPaola, 2009; Saraste, A. and K. Pulkki, 2000). In contrast to its tumor-suppressing effects, autophagy may also help tumor cells survive under hypoxic and nutrient-deprived conditions. Tumorigenic cells with defective apoptosis but functioning autophagy display a survival advantage under ischemic conditions compared with cells with blocked autophagy (Scholzen, T. and J. Gerdes, 2000). Due to the opposing effects of autophagy in tumorigenesis, targeting the autophagic pathway in anticancer therapy may be particularly difficult. For example, inhibition of autophagy increased radiation sensitivity in radioresistant human cancer cell lines in one study (Singh, V.K. et al., 2015; Pang X et al., 2026), whereas another study showed that induction of autophagy radiosensitized prostate cancer cells (Villanueva, A. et al., 2008). Treatment of lymphoma cells with the p53 stimulator tamoxifen led to tumor apoptosis and increased autophagy in surviving cells. Co-treatment with chloroquine enhanced p53-dependent apoptosis and tumor regression by blocking autophagy (Sinha, M. et al., 2012). Therefore, modulation of autophagy as an adjuvant to standard chemotherapy may improve efficacy by shifting the balance toward apoptosis, but this needs to be tumor- and tumor-stage-specific (Waselenko, J.K. et al., 2004). A similar strategy can be employed for the survival of radiation-exposed normal tissues by modulating autophagy levels.

Our 30-day survival study in mice has confirmed that Rap can improve survival following an 8 Gy total-body γ-irradiation (TBI) dose compared with IR alone (Coleman, C.N. et al., 2004). The manifestation of gastrointestinal syndrome, at doses above 6 Gy, is the common cause of death of irradiated animals. Following exposure to ionizing radiation, cells at the base of crypts undergo rapid apoptosis or temporary or permanent senescence, depending on the absorbed radiation dose (Potten, C.S.,1990). This causes malabsorption, electrolyte imbalances, diarrhea, inflammation, infections, weight loss, and ultimately results in mortality (Yuan, R. et al., 2009). Our study showed that a single prophylactic dose of Rap (2 mg /kg body weight) before sublethal irradiation (8 Gy) countered the radiation-induced atrophy of the mucosal layer, decrease in jejunum villi number and cellularity, etc., and resulted in a reduction of apoptosis (Figure 2A, 2B, 5A, and 5B). The findings in this study suggest a potential involvement of autophagy-related processes, as indicated by changes in the autophagy markers LC3 and p62/SQSTM1. However, it should be noted that autophagic flux was not directly assessed in this study (Figure 4A and 4B). Measurements of LC3-II and p62 alone cannot distinguish between enhanced autophagosome formation and impaired lysosomal degradation; the results should be interpreted with caution. Future studies employing flux-based approaches, such as LC3-II turnover assays in the presence of lysosomal inhibitors or ultrastructural analysis, will be important to more precisely define the role of autophagy in this context. In addition, rapamycin also exerts broader biological effects, including metabolic and anti-inflammatory actions. However, the possibility that some of the observed effects may involve additional mTOR-dependent pathways cannot be excluded. Our findings are consistent with a potential contribution of autophagy to the observed protective effects of rapamycin, while recognizing that additional mechanistic studies, such as genetic inhibition of autophagy, would be required to establish an autophagy-specific role.

We observed that radiation exposure (8 Gy) caused severe mucosal layer injury, mainly loss of viable crypt cells and disruption of villus integrity and functionality (Figure 2A). These pathologic developments were significantly reduced by Rap treatment, demonstrating its protective effect against TBI-inflicted GI injury. At an 8 Gy radiation dose, mortality is known to result from the combined effects of gastrointestinal and hematopoietic injury, and recovery of either system may significantly influence survival outcomes. Although our data demonstrate beneficial effects on intestinal morphology and regeneration, the accompanying increase in bone marrow Ki-67 expression suggests that hematopoietic recovery may also contribute to the survival advantage. Consequently, the survival benefit observed in this study likely reflects a systemic response involving multiple radiation-sensitive tissues rather than exclusively enhanced intestinal repair. Furthermore, epithelial regeneration would require compartment-specific quantification of epithelial cells. Future studies should further delineate the relative contributions of gastrointestinal and hematopoietic protection to overall survival. We acknowledge that the exclusive use of female mice in our study is a limitation. Sex can influence multiple biological pathways, including immune and metabolic responses, that may affect the phenotype under study. Although female mice were used throughout the study to ensure experimental uniformity, future investigations that include both male and female animals will be important for evaluating potential sex-dependent differences.

These results demonstrate that a combination treatment with autophagy modulators can be an effective adjuvant strategy, alongside other antioxidants, to augment the IR-induced GI manifestations. An aspect not addressed in this study is the lack of compartment-specific analysis of epithelial cell proliferation. Additionally, in this study, the experimental design evaluates the prophylactic radioprotective effect rather than therapeutic repair following established radiation injury. Forthcoming studies involving post-irradiation dosing will be necessary to determine whether the compound can promote tissue repair after radiation exposure.

## Author Contributions

K.S. conceived the project. M.C. performed most of the experiments and analysis. Technical assistance with flow cytometry measurements and Western blotting was provided by A.S. and K.N. The manuscript was written and edited by M.C. and K.S. All authors reviewed and approved the final version of the manuscript for submission.

## Ethics Statement

The animal study protocol was approved by the Institutional Ethics Committee of the Institute of Nuclear Medicine and Allied Sciences (INMAS) for studies involving animals (protocol code: 8/GO/RBi/S/99/CPCSEA; date of approval: 17.10.2016).

## Data Availability Statement

The original contributions presented in this study are included in the article/supplementary material. Further inquiries can be directed to the corresponding author.

## Acknowledgments

The authors thank Dr. A. K. Singh, Former Director, INMAS, for their constant encouragement. We also thank Dr. Kailash Manda and Dr. Ravi Soni from INMAS for their technical support and valuable suggestions. We thank Dr. N. K. Choudhury for providing access to his fluorescence microscopy facility, Ms. Teena Haritwal, and Dr. Paban Agarawala for histology section preparation.

## Abbreviations

The following abbreviations are used in this manuscript:

ACTB: Actin beta
ATG: Autophagy-related proteins
CQ: Chloroquine
H&E: Hematoxylin and Eosin
IHC: Immunohistochemistry
IR: Ionizing radiation
LC3: Microtubule-associated protein 1 light chain
MTOR: Mechanistic target of rapamycin kinase
PARP1: PolyADP-ribose polymerase 1
PRKAA1: Protein kinase AMP-activated catalytic subunit alpha 1
Rap: Rapamycin
ROS: Reactive oxygen species
TUNEL: Terminal deoxynucleotidyl transferase (TdT) dUTP Nick-End Labeling

## References

1. Amaravadi, R.K., et al., Autophagy inhibition enhances therapy-induced apoptosis in a Myc-induced model of lymphoma. J Clin Invest, 2007. 117(2): p. 326–36.

2. Apel, A., et al., Blocked autophagy sensitizes resistant carcinoma cells to radiation therapy. Cancer Res, 2008. 68(5): p. 1485–94.

3. Ayala, A., M.F. Munoz, and S. Arguelles, Lipid peroxidation: production, metabolism, and signaling mechanisms of malondialdehyde and 4-hydroxy-2-nonenal. Oxid Med Cell Longev, 2014. 2014: p. 360438.

4. Bala, M., et al., Sea Buckthorn Leaf Extract Protects Jejunum and Bone Marrow of (60) Cobalt-Gamma-Irradiated Mice by Regulating Apoptosis and Tissue Regeneration. Evid Based Complement Alternat Med, 2015. 2015: p. 765705.

5. Cao, C., et al., Inhibition of mammalian target of rapamycin or apoptotic pathway induces autophagy and radiosensitizes PTEN null prostate cancer cells. Cancer Res, 2006. 66(20): p. 10040–7.

6. Chaurasia, M., et al., Radiation-induced autophagy: mechanisms and consequences. Free Radic Res, 2016. 50(3): p. 273–90.

7. Verma N, Tiku AB. Role of mTOR pathway in modulation of radiation-induced bystander effects. Int J Radiat Biol. 2022;98(2):173–82.

8. Chaurasia, M., et al., Radiation induces EIF2AK3/PERK and ERN1/IRE1 mediated pro-survival autophagy. Autophagy, 2019. 15(8): p. 1391–1406.

9. Chen, N. and J. Debnath, Autophagy and tumorigenesis. FEBS Lett, 2010. 584(7): p. 1427–35.

10. Chen, Y., M.B. Azad, and S.B. Gibson, Superoxide is the major reactive oxygen species regulating autophagy. Cell Death Differ, 2009. 16(7): p. 1040–52.

11. Coleman, C.N., et al., Medicine. Modulation of radiation injury. Science, 2004. 304(5671): p. 693–4.

12. Eke I, Makinde AY, Aryankalayil MJ, Sandfort V, Palayoor ST, Rath BH, et al. Exploiting Radiation-Induced Signaling to Increase the Susceptibility of Resistant Cancer Cells to Targeted Drugs: AKT and mTOR Inhibitors as an Example. Mol Cancer Ther. 2018;17(2):355–67.

13. Dainiak, N., Hematologic consequences of exposure to ionizing radiation. Exp Hematol, 2002. 30(6): p. 513–28.

14. Darzynkiewicz, Z., D. Galkowski, and H. Zhao, Analysis of apoptosis by cytometry using TUNEL assay. Methods, 2008. 44(3): p. 250–4.

15. Degenhardt, K., et al., Autophagy promotes tumor cell survival and restricts necrosis, inflammation, and tumorigenesis. Cancer Cell, 2006. 10(1): p. 51–64.

16. Dorr, H. and V. Meineke, Acute radiation syndrome caused by accidental radiation exposure - therapeutic principles. BMC Med, 2011. 9: p. 126.

17. Gera, J. and A. Lichtenstein, The mammalian target of rapamycin pathway as a therapeutic target in multiple myeloma. Leuk Lymphoma, 2011. 52(10): p. 1857–66.

18. Homewood, C.A., et al., Lysosomes, pH and the anti-malarial action of chloroquine. Nature, 1972. 235(5332): p. 50–2.

19. Sobecki M, Mrouj K, Camasses A, Parisis N, Nicolas E, Lleres D, et al. The cell proliferation antigen Ki-67 organises heterochromatin. Elife. 2016;5:e13722.

20. Kim WH, Yoo JH, Yoo IK, Kwon CI, Hong SP. Effects of Mesenchymal Stem Cells Treatment on Radiation-Induced Proctitis in Rats. Yonsei Med J. 2023;64(3):167–74.

21. Chen Y, Liu X, Tong Z. Mesenchymal Stem Cells in Radiation-Induced Pulmonary Fibrosis: Future Prospects. Cells. 2022;12(1).

22. Fliedner, T.M., et al., Pathophysiological principles underlying the blood cell concentration responses used to assess the severity of effect after accidental whole-body radiation exposure: an essential basis for an evidence-based clinical triage. Exp Hematol, 2007. 35(4 Suppl 1): p. 8–16.

23. Galluzzi, L., et al., Essential versus accessory aspects of cell death: recommendations of the NCCD 2015. Cell Death Differ, 2015. 22(1): p. 58–73.

24. Ghosh, S.P., et al., Amelioration of radiation-induced hematopoietic and gastrointestinal damage by Ex-RAD(R) in mice. J Radiat Res, 2012. 53(4): p. 526–36.

25. Kim, Y.J., E.H. Kim, and K.B. Hahm, Oxidative stress in inflammation-based gastrointestinal tract diseases: challenges and opportunities. J Gastroenterol Hepatol, 2012. 27(6): p. 1004–10.

26. Hall EJ, Giaccia AJ. eds. Radiobiology for the Radiologist, 6th ed. Philadelphia, PA, USA: Lippincott Williams & Wilkins, (2006).

27. Wu W, Cai Y, Yang Z, Chen M, Hu J, Qu K, et al. Radiation-induced intestinal injury: from molecular mechanisms to clinical translation. Oncol Rev. 2025;19:1613704

28. Kim, Y.C., et al., Bone Marrow Protein Oxidation in Response to Ionizing Radiation in C57BL/6J Mice. Proteomes, 2014. 2(3): p. 291–302.

29. Hacker, G., The morphology of apoptosis. Cell Tissue Res, 2000. 301(1): p. 5–17.

30. Klionsky, D.J., et al., Guidelines for the use and interpretation of assays for monitoring autophagy (4th edition)(1). Autophagy, 2021. 17(1): p. 1–382.

31. Lang, S.A., et al., Mammalian target of rapamycin is activated in human gastric cancer and serves as a target for therapy in an experimental model. Int J Cancer, 2007. 120(8): p. 1803–10.

32. Laplante, M. and D.M. Sabatini, Regulation of mTORC1 and its impact on gene expression at a glance. J Cell Sci, 2013. 126(Pt 8): p. 1713–9.

33. Levine, B., Autophagy and cancer. Nature, 2007. 446(7137): p. 745–747.

34. Levine, B. and G. Kroemer, Autophagy in the pathogenesis of disease. Cell, 2008. 132(1): p. 27–42.

35. MacNaughton, W.K., Review article: new insights into the pathogenesis of radiation-induced intestinal dysfunction. Aliment Pharmacol Ther, 2000. 14(5): p. 523–8.

36. Marklund S, Marklund G. Involvement of the superoxide anion radical in the autoxidation of pyrogallol and a convenient assay for superoxide dismutase. Eur J Biochem. 1974 Sep 16;47(3):469–74. doi: 10.1111/j.1432-1033.1974.tb03714.x. PMID: 4215654.

37. Measey, T.J., et al., Pilot Study of Radiation-induced Gastrointestinal Injury in a Hemi-body Shielded Gottingen Minipig Model. Health Phys, 2018. 114(1): p. 43–57.

38. Meistrich, M.L. and M. Kangasniemi, Hormone treatment after irradiation stimulates recovery of rat spermatogenesis from surviving spermatogonia. J Androl, 1997. 18(1): p. 80–7.

39. Meng, L.H. and X.F. Zheng, Toward rapamycin analog (rapalog)-based precision cancer therapy. Acta Pharmacol Sin, 2015. 36(10): p. 1163–9.

40. Moron MS, Depierre JW, Mannervik B. Levels of glutathione, glutathione reductase and glutathione S-transferase activities in rat lung and liver. Biochim Biophys Acta. 1979 Jan 4;582(1):67–78. doi: 10.1016/0304-4165(79)90289-7. PMID: 760819.

41. Njie-Mbye, Y.F., et al., Lipid peroxidation: pathophysiological and pharmacological implications in the eye. Front Physiol, 2013. 4: p. 366.

42. White, E. and R.S. DiPaola, The double-edged sword of autophagy modulation in cancer. Clin Cancer Res, 2009. 15(17): p. 5308–16.

43. Saraste, A. and K. Pulkki, Morphologic and biochemical hallmarks of apoptosis. Cardiovasc Res, 2000. 45(3): p. 528–37.

44. Scholzen, T. and J. Gerdes, The Ki-67 protein: from the known and the unknown. J Cell Physiol, 2000. 182(3): p. 311–22.

45. Singh, V.K., et al., Animal models for acute radiation syndrome drug discovery. Expert Opin Drug Discov, 2015. 10(5): p. 497–517.

46. Pang X, Liu H, Long Y, Wang H. *mTOR in radiotherapy of lung cancer: Mechanisms of radiation resistance and therapeutic implications* (Review). Int J Oncol. 2026;68(2).

47. Villanueva, A., et al., Pivotal role of mTOR signaling in hepatocellular carcinoma. Gastroenterology, 2008. 135(6): p. 1972-83, 1983 e1-11.

48. Sinha, M., et al., Amelioration of ionizing radiation induced lipid peroxidation in mouse liver by Moringa oleifera Lam. leaf extract. Indian J Exp Biol, 2012. 50(3): p. 209–15.

49. Waselenko, J.K., et al., Medical management of the acute radiation syndrome: recommendations of the Strategic National Stockpile Radiation Working Group. Ann Intern Med, 2004. 140(12): p. 1037–51.

50. Potten, C.S., A comprehensive study of the radiobiological response of the murine (BDF1) small intestine. Int J Radiat Biol, 1990. 58(6): p. 925–73.

51. Varshney R, Kale RK. Effects of calmodulin antagonists on radiation-induced lipid peroxidation in microsomes. Int J Radiat Biol. 1990 Nov;58(5):733–43. doi: 10.1080/09553009014552121. PMID: 1977818.

52. Yuan, R., et al., Targeting tumorigenesis: development and use of mTOR inhibitors in cancer therapy. J Hematol Oncol, 2009. 2: p. 45.

