## Supplementary Figure S1 for "Targeting Autophagy Accelerates Intestinal Repair after Acute Ionizing Radiation"

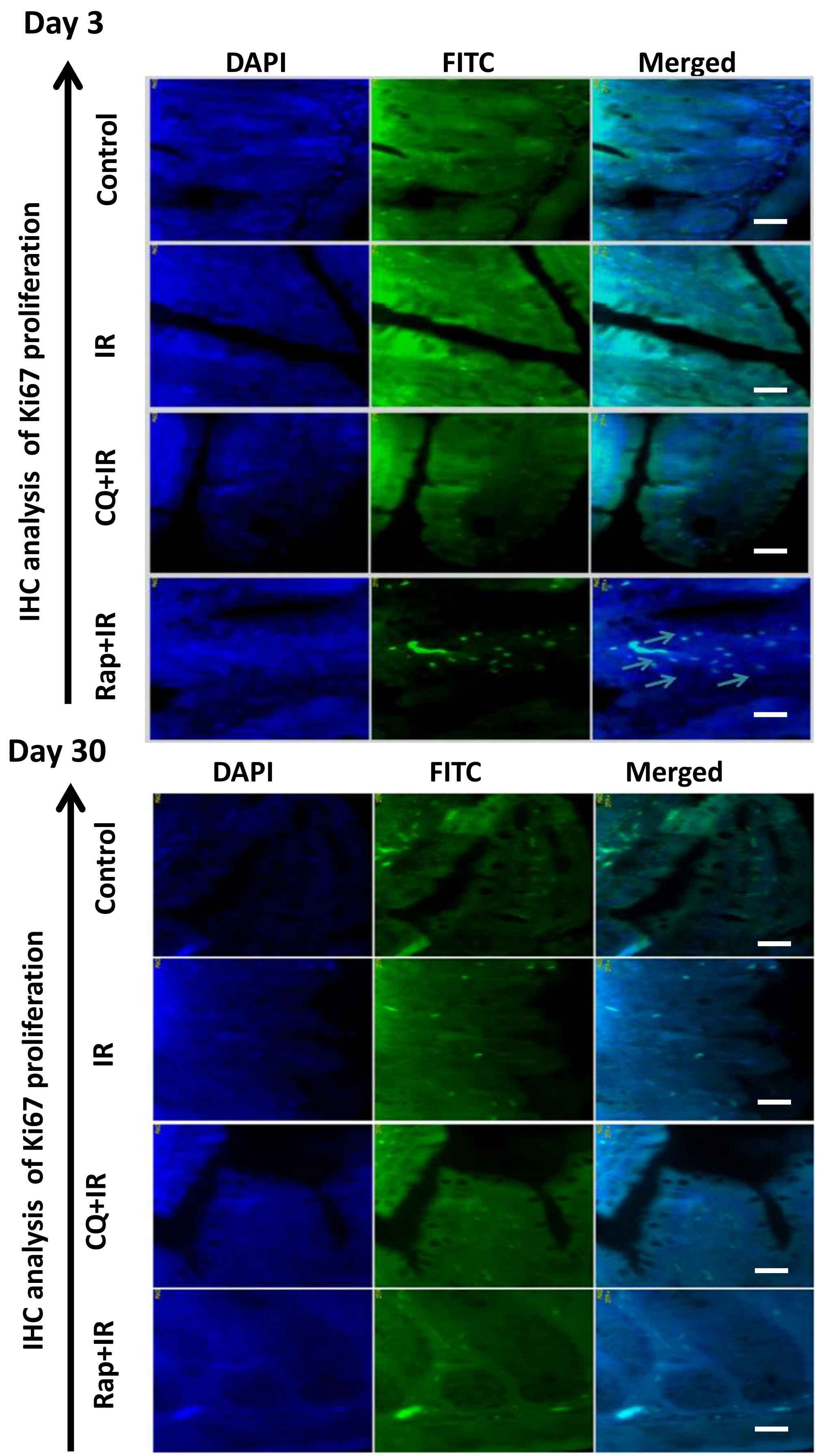


**Figure S1.** Representative visualization of fluorescence-based immunohistochemistry (IHC-F) of Ki-67-positive cell proliferation in the presence of autophagy modifier drugs on day 3 and day 30 post-exposure.
