## Supplementary material for "Targeting Autophagy Accelerates Intestinal Repair after Acute Ionizing Radiation": supplemetary figure S2

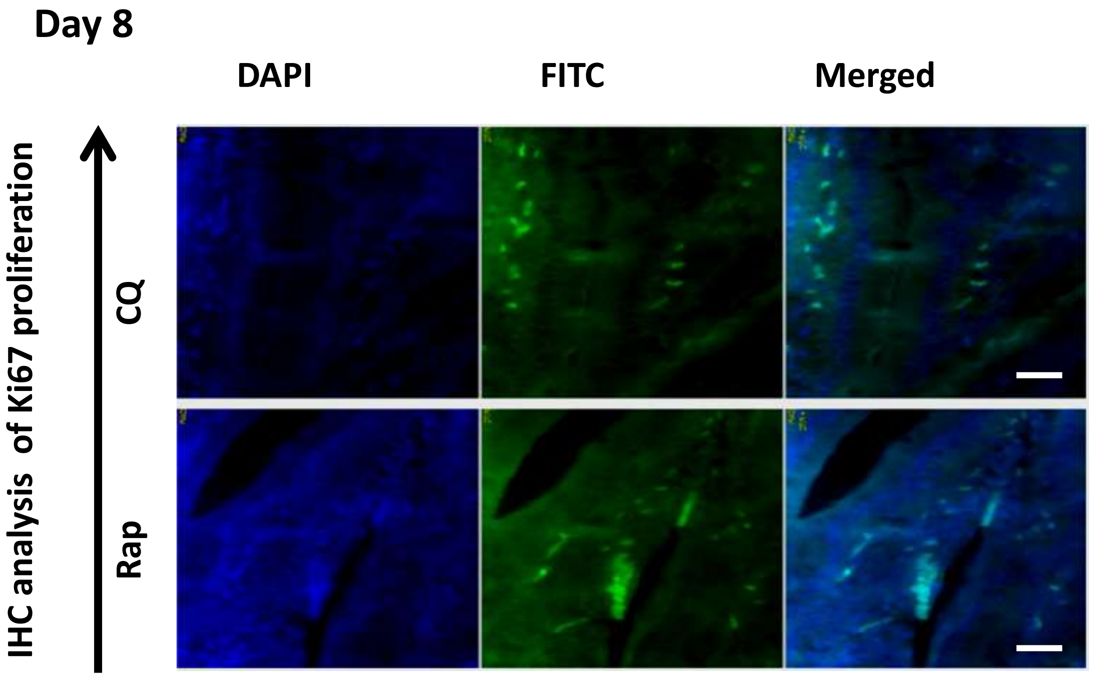


**Figure S2.** Representative visualization of fluorescence-based IHC-F analysis of Ki-67-staining in intestinal tissue in the presence of drug-alone groups, i.e., Rap and CQ alone, on day 8 post-exposure.
